# On the regulatory evolution of new genes throughout their life history

**DOI:** 10.1101/276667

**Authors:** Jia-yu Zhang, Qi Zhou

## Abstract

Every gene has a birthplace and an age, i.e., a cis-regulatory environment and an evolution lifespan since its origination, yet how gene’s evolution trajectory is shaped by the two remains unclear. Here we address this basic question by comparing phylogenetically dated new genes of different ages among Drosophila and vertebrate species. For both, we find a clear ‘out of testis’ transition from the testis-specific young genes to the broadly expressed old housekeeping genes. Particularly, many new genes have evolved specific activation at maternal-zygotic transition, or distinctive spatiotemporal embryonic expression patterns from the parental genes. We uncover an age-dependent gain/loss of active/repressive histone modifications and cis-regulatory elements, with variations between species and between somatic/germline tissues, which together underpin the stepwise acquisition of novel and important function by new genes. These results illuminate the general evolution trajectory and the underlying regulatory mechanisms of genes throughout their life history.

## Introduction

The great disparity of gene numbers between species indicates that gain and loss of genes is a fundamental evolution process. Since the report of the first new gene *jingwei* over two decades ago(Long & Langley, 1993), numerous genome-wide and case studies have now demonstrated that origination of functional new genes is one of the main drivers underlying phenotypic innovation(S. Chen, Krinsky, & Long, 2013; Kaessmann, 2010). The emergence of *jingwei* represents a paradigm rather than an anecdote of new gene evolution: both DNA- and RNA-mediated (retroposition) gene duplications from different parental genes have contributed to the formation of the new chimeric gene structure of *jingwei*, which acquired a new expression pattern specifically in testis compared to its *Adh* ancestor. Later inspection of multiple Drosophila genomes showed that gene duplication accounts for about 80% of species or lineage specific new genes(Zhou et al., 2008). This conforms to Ohno’s hypothesis that gene duplication is the primary source of new genes(Ohno, 1970). In addition, at least 30% of Drosophila new genes(Zhou et al., 2008), or 50% of *Caenorhabditis elegans* new genes(Katju & Lynch, 2006) have been found to incorporate various genomic resources (e.g., partial coding sequences of another gene, or transposable elements) to form a chimeric structure by exon shuffling, potentially facilitating functional innovation. An unexpected finding from genome scans of a broad range of species including yeast(Carvunis et al., 2012), Drosophila(Zhao, Saelao, Jones, & Begun, 2014) and human(Knowles & McLysaght, 2009; Ruiz-Orera et al., 2015; Wu, Irwin, & Zhang, 2011) is that *de novo* origination from non-coding sequences has a substantial contribution to new gene origination. Many nascent *de novo* genes, as well as species-specific gene duplicates are more likely to be still segregating within populations and subjected to random loss than those ‘older’ new genes that have become fixed in populations at an earlier time point and are shared by multiple species(Palmieri, Kosiol, & Schlotterer, 2014; Zhao et al., 2014; Zhou et al., 2008). Similar to *jingwei*, many *de novo* genes and new gene duplicates have been found to become predominantly or exclusively expressed in testis(Carelli et al., 2016; Guschanski, Warnefors, & Kaessmann, 2017; Luis Villanueva-Canas et al., 2017). Functional disruption showed some Drosophila new genes acquired novel function that is either involved in spermatogenesis(Kondo et al., 2017)(e.g., *nsr* (Ding et al., 2010) gene that originated about 6 million years ago, MYA), or associated with male mating behavior (e.g., *sphinx* (Dai et al., 2008) that originated 3 MYA). A striking case is *Umbrea*, an older (15 MYA) new gene that gradually evolved essential centromeric function in comparison to its heterochromatin-binding parental gene HP1B(Ross et al., 2013). These case studies suggested that new genes frequently undergo neo-functionalization, and their population dynamics and novel function is characterized by their age.

Understanding functional evolution of new genes in the context of their ages is critical for illuminating genes’ dynamic life history in general(Betran, 2015; Carelli et al., 2016). Although it is difficult to reconstruct gene’s evolution trajectory, valuable insights have been gained by comparing genes of different ages(Carelli et al., 2016; Guschanski et al., 2017). This is on one hand facilitated by the ongoing effort of functional disruption of identified Drosophila new genes using RNAi or CRISPR/Cas-9 technique(Sidi Chen, Zhang, & Long, 2010; Kondo et al., 2017), and also by the recent development of next-generation sequencing. Transcriptome comparison of multiple Drosophila and mammalian adult tissues suggested that younger new gene duplicates, particularly retrogenes are more prone to have a testis-specific expression; while the older ones are more often ubiquitously expressed or specifically expressed in other somatic tissues(Assis & Bachtrog, 2013; Carelli et al., 2016; Guschanski et al., 2017). This has led to the ‘out of the testis’ hypothesis on the emergence of new genes: it postulates that the permissive chromatin environment of testis provides a haven for nascent genes from natural selection against deleterious effects of the redundant gene dosage when they are born(Kaessmann, 2010). These young genes maybe later driven to fixation by intensive sexual selection in testis, or acquisition of novel function beyond the testis by forming new gene structures and/or recruiting new regulatory elements. Such a dynamic life history of new genes is also reflected by the gradually increased connectivity of gene interactions from young genes usually located at the periphery of the network to old genes as an essential hub(W. Zhang, Landback, Gschwend, Shen, & Long, 2015). Overall, most contemporary genome-wide portraits of new genes take advantage of transcriptome data, which is the output of complex coordinated regulation involving cis-regulatory elements (CREs: promoter, enhancer etc.) and local epigenomic configuration.

However, little is known about the regulatory mechanisms underlying how a new gene evolves a divergent expression pattern from its ancestor at the genome-wide level. This is because components and principles of transcriptional regulation have not been systematically dissected only until very recently by many consortium projects (e.g., ENCODE and modENCODE)(E. P. Consortium, 2012; G. Consortium et al., 2017; m. Consortium et al., 2010). This question is key to understanding how a new gene can avoid becoming a pseudogene as presumed by the classic model(Ohno, 1970). A new gene can either evolve new expression, i.e., undergo neo-functionalization by recruiting novel cis-regulatory elements, and/or translocating to a new epigenomic environment as often occurred with retrogenes. Alternatively, a new gene can partition the ancestral expression pattern with its parental gene through complementary degenerative mutations in the regulatory region (sub-functionalization)(Lynch & Force, 2000). It is now well established that the epigenomic landscape is shaped by dynamic DNA methylation and various histone modifications. Active and repressive chromatin marks, such as histone H3 lysine 4 trimethylation (H3K4me3), H3K36me3, and H3K27me3, H3K9me3 etc. synergistically or antagonistically bind together to genic or CRE regions to impact the transcription level. In this work, we seek to address the regulatory mechanisms of new gene evolution by analysing a total of 83 transcriptomic and 281 epigenomic datasets across a broad range of tissues and developmental stages of Drosophila and human (**Supplementary Fig. 1**). We used an updated dataset of new genes of Drosophila and vertebrates, and paid special attention to bulk and single-cell RNA-seq (scRNA-seq) data during early development, of which little is known regarding new genes’ functional role. By cross investigation of the epigenomic and CRE profiles of new genes with the context of their origination mechanisms and ages, we unveiled a complete landscape of dynamic regulatory changes throughout a new gene’s life history.

**Figure. 1:**
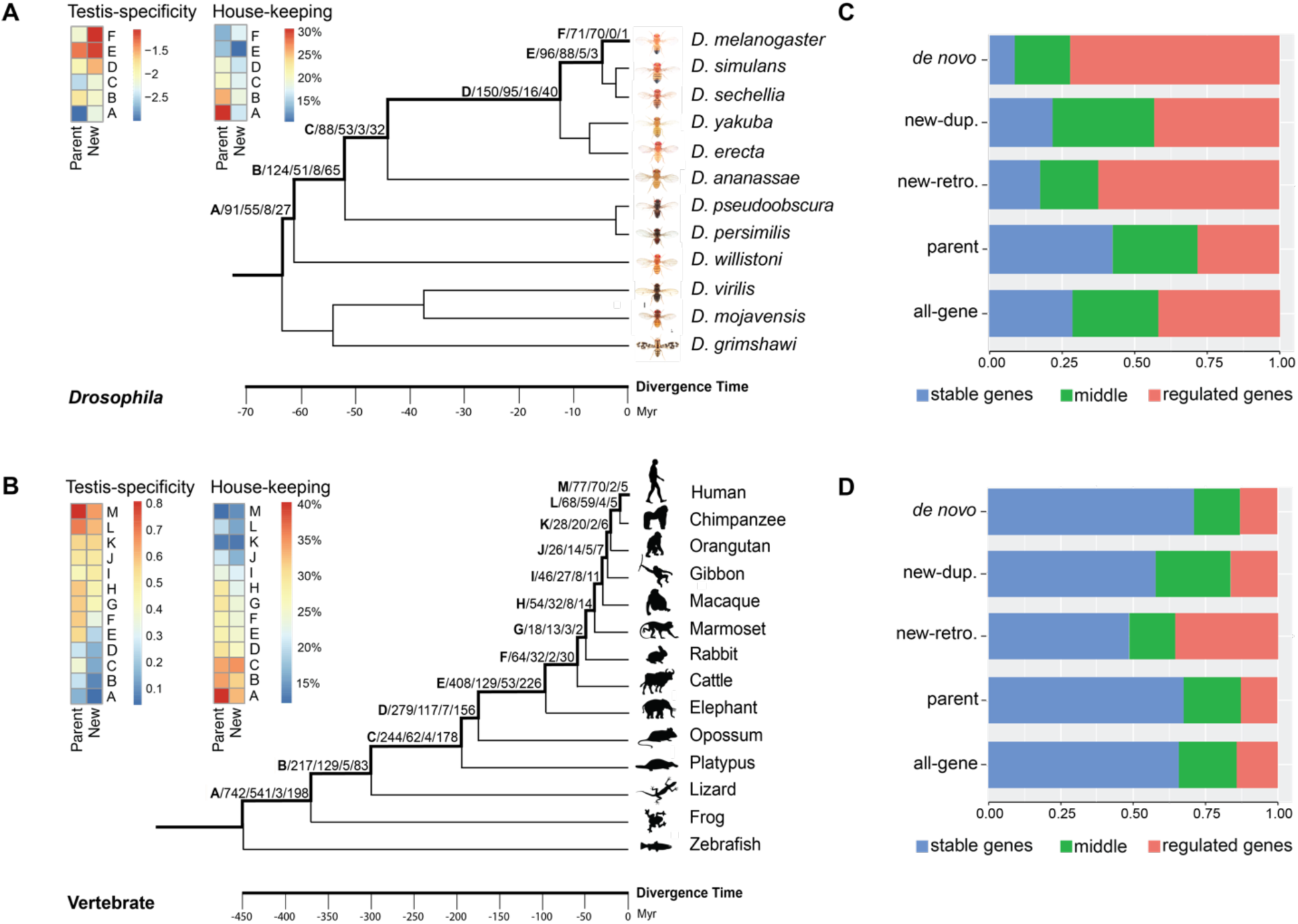
Drosophila and vertebrate new genes and their expression patterns. (A-B) Different numbers of new genes divided by their age and origination mechanisms. Each age group is designated by an ordered letter from A to F/M shown on each phylogenetic branch. And each group of genes is further divided as total number of new genes, number of DNA-based duplicated genes, retrogenes, and *de novo* genes, separated by slash. We also showed along the age groups the change of testis-specificity, measured by the log2 based ratio of expression level of testis vs. whole male body; and housekeeping gene index, measured by the percentage of tissues/stages which show robust expression. Median values of testis specificity and housekeeping gene index of each age group is shown. The Drosophila pictures are from Nicolas Gompel. (C-D) Regulated genes of human and Drosophila, divided by different origination mechanisms. We defined the regulated genes based on the coefficient of expression (CV) level variation across different tissues. CVs of *Drosophila* were derived from(Perez-Lluch et al., 2015) and those of human were calculated from GTEx dataset (https://www.gtexportal.org/)(G. Consortium et al., 2017).

## Results

### New genes are becoming out of testis by age

We acquired a high-confidence dataset of new genes following the published pipeline(Sidi Chen et al., 2010; Y. E. Zhang, Vibranovski, Landback, Marais, & Long, 2010) with the updated genomes of 12 Drosophila species (metazoa release 25) and 14 vertebrate species (Ensembl v73) (Fig. 1). In brief, we used whole-genome syntenic alignments to inspect the phylogenetic distribution of orthologous genes. We identified species or lineage specific new genes and inferred their age by parsimony based on their presence/absence of orthologs in multiple outgroups. We also inferred their origination mechanisms as gene duplication (specifically referring to DNA-based duplication hereafter), retroposition and *de novo* origination based on each category of a gene’s specific feature (e.g., absence of introns in retrogenes, see **Methods**). In total, we annotated 585 Drosophila and 3056 vertebrate new genes. A major technical challenge for any new gene analyses is that except for *de novo* genes, a large part of the sequences is identical between the new and parental gene pair, which confounds the comparison of their gene expression level or histone modification. To overcome this, we harnessed the sequence divergence sites between the two, and only counted reads that span such informative sites throughout this work. Three lines of evidence convinced us that this subset of sequences per gene is able to give us robust and specific estimation of the level of transcription and histone modification: first, the SNP density, as a reflection of divergence level between the gene pair expectedly increases by new genes’ age (**Supplementary Fig. 1**). We found a substantial number of informative sites (median value 10-30 per 100 bp) even between the youngest group of new genes and their parental genes, given a read length of over 100 bp in most of the analysed datasets. Second, we observed a significant (*P*<0.05, t-test) positive/negative correlation between the normalized expression level vs. respective active/repressive histone modification level (**Supplementary Fig. 1**) across all the inspected stages and tissues, based only on informative sites. Third, the proportions of genes showing pronounced histone modifications (‘bound’ genes), defined by informative sites, increased gradually with the developmental progress of Drosophila (**Supplementary Fig. 1**). This pattern is consistent with that reported by modENCODE which used the entire gene or promoter region(Kharchenko et al., 2011).

**Figure. 2:**
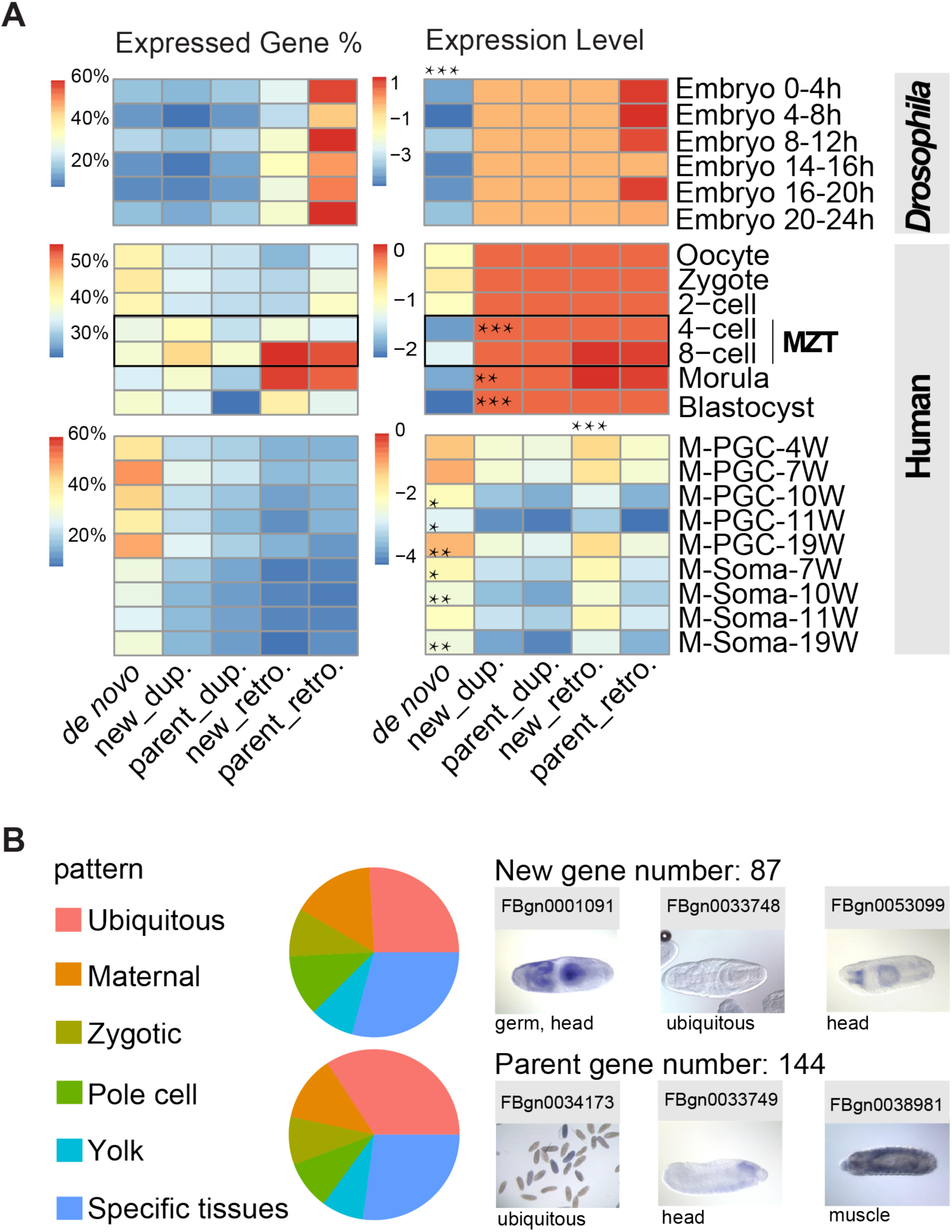
Dynamic expression of new genes in early embryos. (A) We showed the percentage of active genes (left columns), and the median values of normalized gene expression levels (right columns) across different embryonic stages of *Drosophila* (with bulk RNA-seq) and human (with single RNA-seq), ordered by the developmental course. We also compared the new genes vs. parental genes, and *de novo* genes vs. the genome-wide average for their expression levels and show the level of significance of t-test (*: *P* < 0.05; **: *P* < 0.01; ***: *P* <0.001). If the entire column is significant, we show the asterisks above the column (e.g., human retrogenes). M-PGC-4W: male primordial germ cell of the 4 weeks’ stage. (B) Divergent subcellular localization between new and parental genes in Drosophila embryos. Pie charts showed the proportion of different expressed locations of new genes and parent genes, with examples of divergent expression patterns between new and parental genes.

**Figure. 3:**
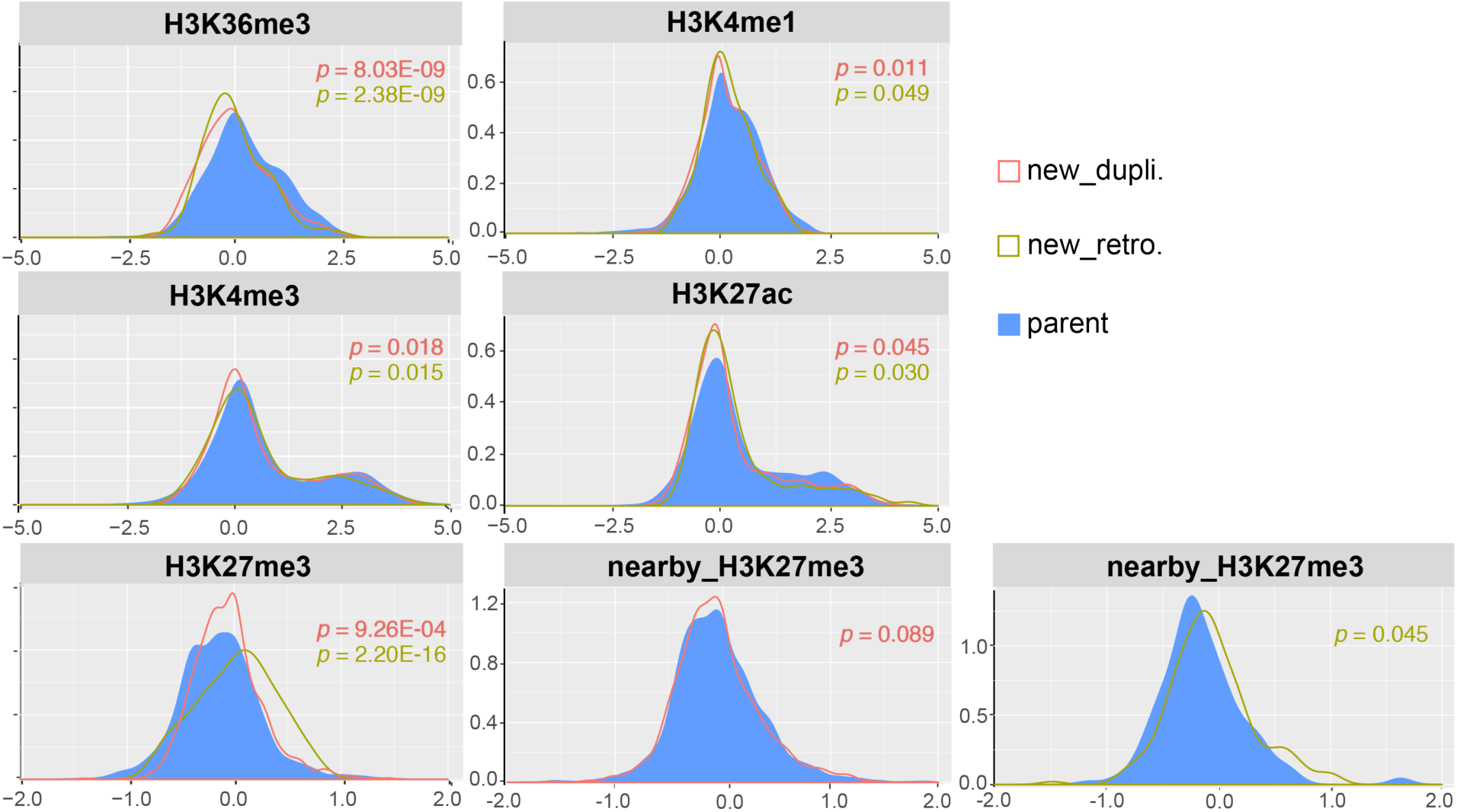
New genes and parental genes have diverged for their chromatin state. We showed the distributions of normalized histone modification levels measured by log2 ChIP vs. input ratio, spanning the gene body or promoter regions of new gene (different colors of lines) vs. parental gene (blue area), and their surrounding genes (‘nearby’ profiles). We also showed the *P*-values of t-tests comparing the new genes vs. parental genes.

**Figure. 4:**
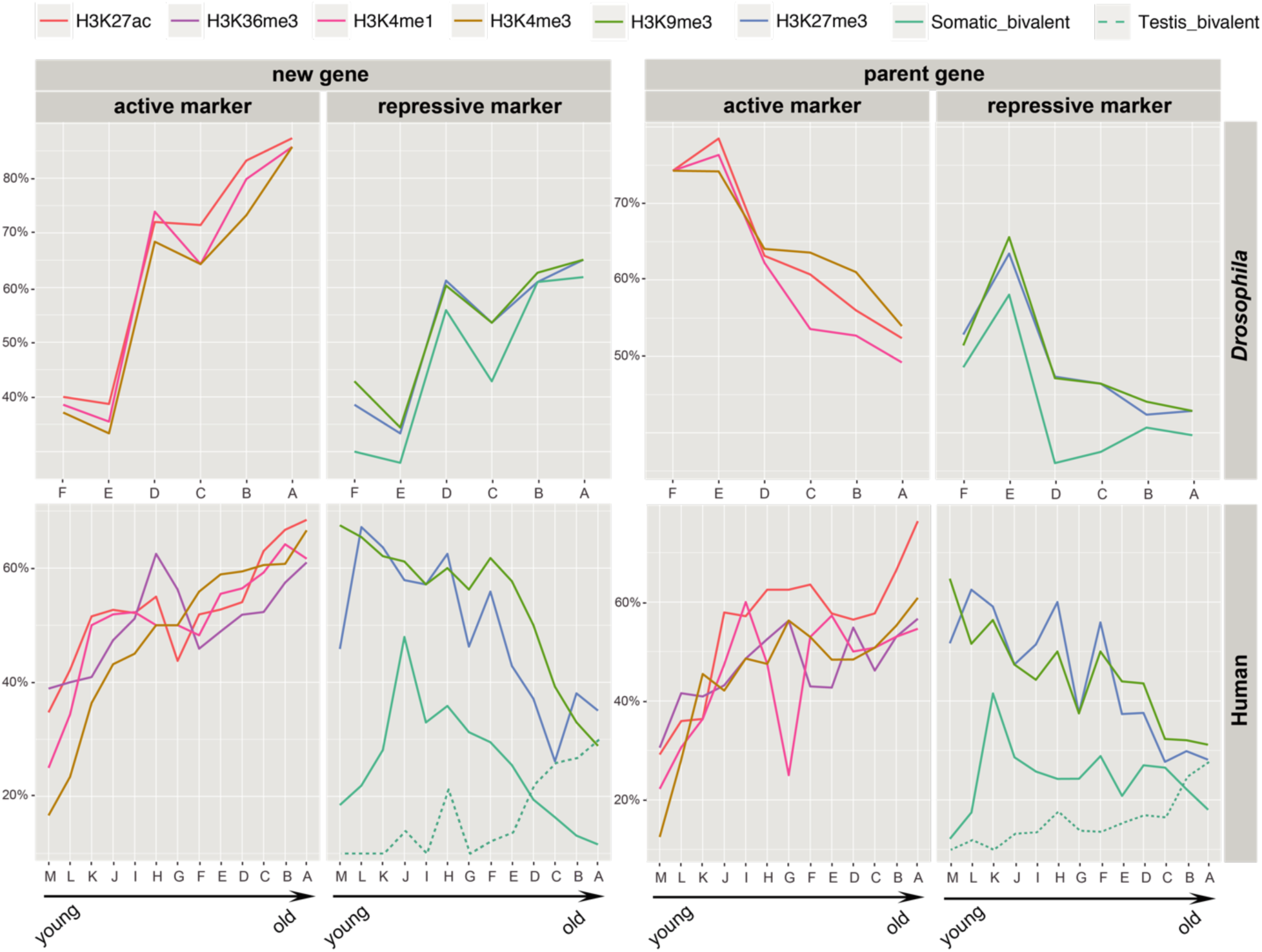
Age-dependent change of histone modifications. We show here the percentage of bound genes by respective histone modifications across the ordered age groups. The data of human testis bivalent genes are derived from(Lesch et al., 2016) shown as dotted line. For the solid lines, we used the ChIP-seq data of second instar larvae for *Drosophila*, and adrenal gland adult male tissue for human. The patterns of other Drosophila or human tissues are consistent and shown in Supplementary Fig. 18.

To reveal the global transcriptome change of new genes by age, we focused on two of their features: testis specificity, calculated as the ratio of gene expression level in testis vs. that of the whole male body; and expression breadth, measured by the percentage of tissues/stages with active expression of the focal gene detected among all examined samples, as an indication for housekeeping genes. It is clear that in both Drosophila and human, younger new genes are less likely to be housekeeping genes and more likely to be testis-specific, supporting the ‘out of testis’ hypothesis (Fig. 1A-B). Interestingly, the same trend has also been found for parental genes, suggesting strong selection against duplication of housekeeping genes has also contributed to this pattern. As the expression breadth *per se* does not reflect the degree of expression level differences between tissues/stages, we further calculated the coefficients of variation (CV)(Perez-Lluch et al., 2015) for gene expression levels across all the analysed samples. A gene that shows highly variable spatial/temporal gene expression pattern, i.e., a high CV value, is defined as a ‘regulated’ gene based on the distribution of genome-wide CV values (**Supplementary Fig. 1**), otherwise it is defined as a ‘stable’ gene. We consistently found new genes, particularly retrogenes and Drosophila *de novo* genes have a larger proportion of regulated genes than the parental genes or all the genes throughout the genome (Fig. 1C-D). This can be explained by retrogenes and *de novo* genes being more likely to recruit novel regulatory elements than are new genes generated by gene duplication.

**Figure. 5:**
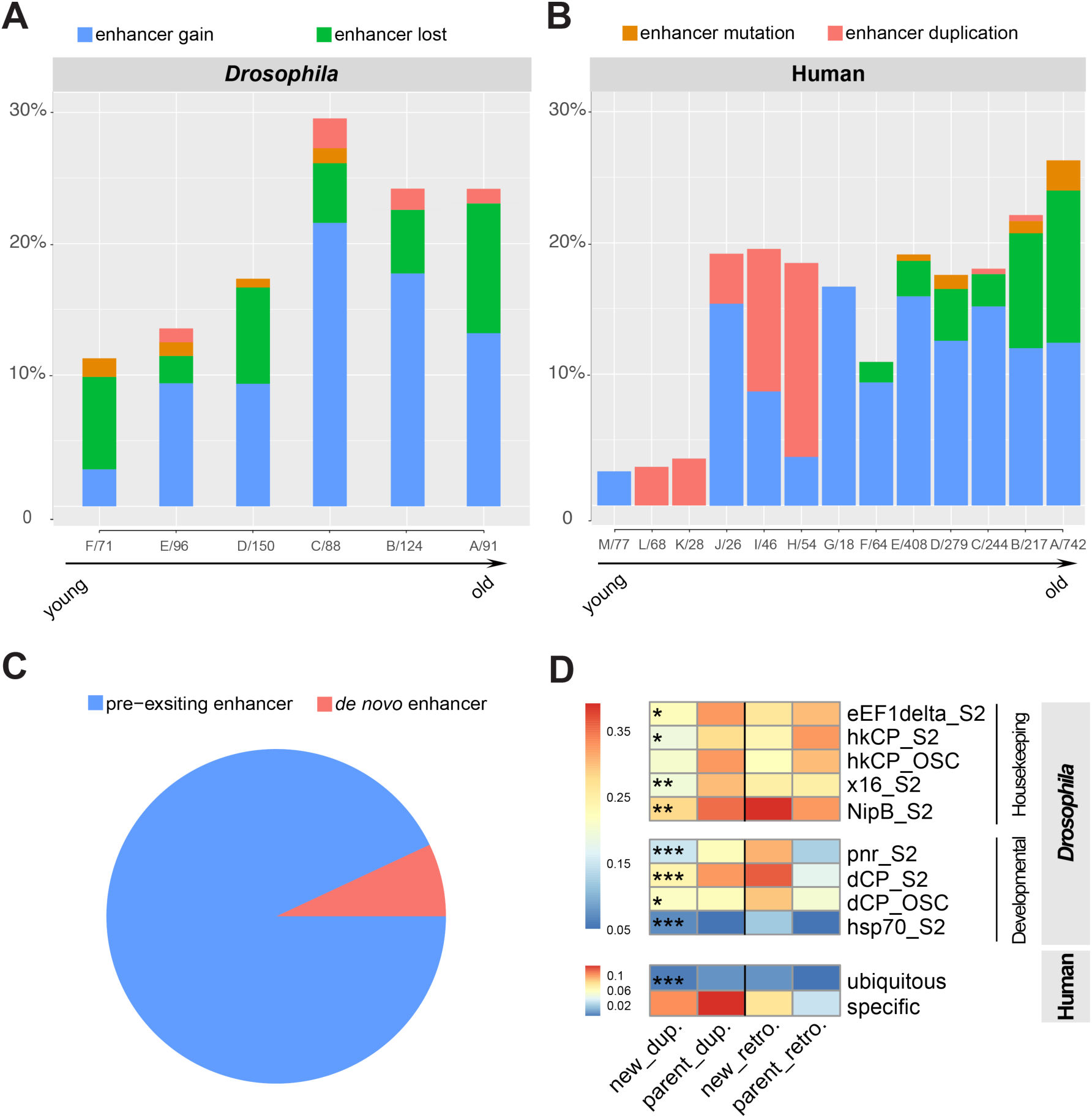
New genes and parental genes have diverged for their CREs. (A-B) We compared the enhancers between new and parental genes in Drosophila and human, based on the latest STARR-seq or CAGE annotations. The bar plot here shows the proportion of new genes at each age group that have undergone turnovers of enhancers. (C) The pie chart shows the source of enhancer gains in Drosophila new genes, either from a pre-existing enhancer, as defined by the presence of an orthologous enhancer in the outgroup species, or a *de novo* enhancer if absent. (D) We compared the numbers of different types of enhancers between new and parental genes, and showed here the average numbers (enhancers per gene) for each category. Each row represents one type of Drosophila developmental or housekeeping gene enhancers identified from(Zabidi et al., 2015), and human ubiquitous or specific enhancers are derived from(Andersson et al., 2014).

### Dynamic expression of new genes during early development

Much effort has been invested in examining new genes’ expression in adult tissues, yet little is known about their functional contribution during the embryonic developmental process. The only work that has been done, at a much broader evolutionary time scale than this work, found that in zebrafish and Drosophila, younger genes are enriched in early and late embryonic stages, while the old genes are enriched at mid-embryonic (‘phylotypic’) stage(Domazet-Loso & Tautz, 2010). This supports a developmental ‘hourglass’ model, where strong constraints on developmental regulation act at the phylotypic stage(Kalinka et al., 2010). Here we scrutinized both the expression level and spatiotemporal information of new genes in early embryos, by analysing their scRNA-seq and RNA fluorescence *in situ* hybridization (RNA-FISH) data. We found in both human and Drosophila, new genes except for Drosophila *de novo* genes show a similarly robust expression level as parental genes throughout embryonic stages, suggesting some of them have evolved important developmental function (Fig. 2A). Human retrogenes and *de novo* genes are expressed at a significantly higher level (*P*<0.05, t-test and Wilcoxon test) respectively than their parental copies and genome-wide average since 4 weeks until 19 weeks of embryonic development, in both germline and somatic cells, in both sexes (**Supplementary Fig. 6)**. We found a burst of expressed retrogenes as well as their expression level, but not in new gene duplicates, from human 4-cell stage to 8-cell stage, in contrast to a decrease of both for *de novo* genes from 2-cell stage until 8-cell stage. This is particularly interesting as the major maternal-zygotic transition (MZT) occurs during this time window(Braude, Bolton, & Moore, 1988). It implies that many retrogenes are involved in MZT, and that although many *de novo* genes are originally deposited as maternal transcripts, they become degraded since the 2-cell stage. Indeed, we found that the zygotically activated retrogenes were enriched for functional categories of ‘RNA-binding’, ‘RNA recognition’ and ‘protein transport’ (**Supplementary Fig. 7**), which might participate in the post-transcriptional control as shown by many genes at Drosophila MZT(Sysoev et al., 2016).

Besides the expression level, we also compared the spatial patterns of embryonic expression between new vs. parental genes of Drosophila, whose different subcellular localizations directly indicate their divergent function. Among the 144 parental genes and 87 new genes with RNA-FISH data available from either BDGP(Tomancak et al., 2002) or Fly-FISH(Lécuyer et al., 2007) database, we found that parental genes are more likely to be ubiquitously expressed (Fig. 2B, 34.25% vs. 25.95% of new genes, *P*=0.0239, Fisher’s exact test) while new genes tend to be expressed within specific tissues (29.20% vs. 27.09% of parental genes, *P*=0.5412 Fisher’s exact test) in embryos. All the 11 *de novo* genes with annotated patterns are all maternal transcripts that become degraded during later development. This is consistent with what was found for human *de novo* genes (Fig. 2A). Among the 40 gene pairs with RNA-FISH data available for both parental and new genes, none of the pairs show exactly the same subcellular localization and expressed time window between the two, providing strong evidence for significant functional divergence between the parental and new genes in early embryos. Specifically, 9 gene pairs (e.g., FBgn0034173-FBgn0001091, Fig. 2B, **Supplementary Fig. 8**) show ubiquitous expression throughout all the investigated stages for the parental gene, but a specific expression pattern for the new gene. 4 pairs show the opposite pattern. These cases are strong candidates for neo-functionalization of new genes (**Supplementary Fig. 9**). There are 27 gene pairs with parental and new genes expressed in different tissues (e.g., FBgn0038901-FBgn0053099, Fig. 2B, **Supplementary Fig. 10**), which can be candidates for sub-functionalization.

### Age-dependent evolution of chromatin state

As many more genes become regulated by histone modifications beyond embryonic stages (**Supplementary Fig. 1**), we further examined the expression patterns of new genes among all the tissues, divided by different age groups. We found human but not Drosophila genes show an age-dependent change of expression: not only new genes but also parental genes become more likely to be expressed in older age groups across all the examined human tissues (**Supplementary Fig. 11-12**). The pattern of *de novo* genes is not as clear, which could be influenced by a smaller number of genes at each age group. Correspondingly, there are gradually less putative pseudogenes, defined as those without robust gene expression throughout all the examined tissues or stages, among older age groups of new genes (**Supplementary Table 1**). This suggested that the tendency to become more active and functionally important is a general age-related feature of genes, not just species or lineage-specific new genes.

To uncover the regulatory mechanisms underlying such an age-dependent expression pattern, we inspected 14 Drosophila and 7 human histone modification markers and first compared their binding patterns between new and parental genes. We focused on four active markers H3K4me3, H3K4me1, H3K27ac, H3K36me3, which are strongly associated with active transcription, and promoter, enhancer or exonic regions; and two repressive markers H3K27me3, H3K9me3 which are associated with gene silencing (**Supplementary Fig. 1**). They were chosen because they are among the best-known for their functional association and most broadly studied across almost all tissues and developmental stages in both human and Drosophila (**Supplementary Fig. 1**)(Ho et al., 2014). We found that in both species, and throughout most stages/tissues, new genes exhibit a significantly (Wilcoxon test, *P*<0.05) lower level of all active histone modifications and RNA Polymerase II binding at promoter (for H3K4me1/3, H3K27ac) or entire gene regions (for H3K36me3), but a higher level of facultative heterochromatin modification H3K27me3 than their parental genes. No significant difference has been observed between new and parental genes for the constitutive heterochromatin modification H3K9me3 (Fig. 3, **Supplementary Fig. 13-14**), which is usually associated with transposable elements. A key distinction between the two repressive markers is that H3K27me3 is strongly associated with spatiotemporal regulation of gene expression, thus more dynamic in its silencing function. In particular, H3K27me3 may form a ‘bivalent’ domain together with H3K4me3 to maintain the influenced gene in a poised chromatin state for later activation of lineage-specific expression. These patterns overall account for a generally lower percentage of expressed genes (**Supplementary Fig. 11**) or housekeeping genes, but a higher percentage of regulated genes, especially retrogenes (Fig. 1) among the new genes comparing to the parental genes. Indeed, we found that retrogenes show an even higher level of H3K27me3 modification than other new genes produced by DNA-based duplication (Fig. 3). This is probably because many retrogenes have translocated into a pre-existing H3K27me3 domain as indicated by their surrounding genes: we found that up- and down-stream genes of new retrogenes also show a significantly (*P*<0.05, Wilcoxon test) higher level of H3K27me3 modification than those surrounding the parental genes, but the pattern of DNA-based duplicated genes is less pronounced or not as consistent as retrogenes across different tissues or stages (Fig. 3, **Supplementary Fig. 15**). This does not indicate retrogenes are more likely to be silenced pseudogenes, as we found that retrogenes show a significantly higher proportion of ‘bivalent’ genes, defined as those bound by both H3K27me3 and H3K4me3 markers, than their parental genes during larvae stages of Drosophila, and in some specific tissues of human (e.g., kidney, **Supplementary Fig. 16-17**).

We further uncovered that both new and parental genes exhibit an age-dependent change of chromatin states, with different trajectories between somatic and germline tissues, and also between human and Drosophila, the latter of which accounts for their presence or absence of age-dependent gene expression pattern. For Drosophila, more new genes become bound by both active and repressive histone marks, and particularly there is a higher proportion of bivalent genes in older age groups. In contrast, their parental genes show an opposite trend. These patterns have been observed from the late stage embryos until the second instar larvae, while other developmental stages have too few bound genes (less than 20% per age groups) to show the pattern (Fig. 4, **Supplementary Fig. 18**). For human, parental and new genes show a similar pattern to each other: there are generally more genes bound by active marks, but fewer genes bound by repressive marks by age, which together result in more active genes in older age groups (**Supplementary Fig. 11**). A similar pattern of active or repressive marks has also been observed for human new and parental genes when we compared the levels of histone modifications among different age groups. The patterns are generally consistent among the investigated somatic tissues or stages, but more pronounced in adult tissues (**Supplementary Fig. 18-19**). Bivalent genes show a more complex pattern along the age groups, and between somatic and germline tissues. There is a burst of bivalent new genes at the ancestor of apes (age group J in **Fig.1, 4**), after which its proportion reduces by age in somatic tissues. While in germline, there is an intriguing ‘into testis’ pattern where the proportion of bivalent genes(Lesch, Silber, McCarrey, & Page, 2016) increases by the age of genes. The opposite trajectory of bivalent genes between somatic and germline tissues makes sense in the light of segregation of novel cell-differentiation related functions of new genes in either type of tissues. These results collectively indicated that it takes young genes some time to evolve active histone modifications, while in human but not in Drosophila, repressive histone modifications have further contributed to the silencing of young genes. They also suggested that strong selection against redundant gene dosage, especially for robustly expressed genes, when a nascent gene copy is born: in human, parental genes of younger new genes tend to be lowly expressed genes with few active marks and many repressive marks; and in Drosophila, while actively expressed parental genes do give birth to new genes of young age, the new genes generally tend to lack active histone modifications to drive robust expression (**Supplementary Fig. 11, 18**).

### New genes and parental genes have become divergent for their CREs

Gene expression is coordinately regulated by epigenomic configuration and CREs. The differential bindings of histone modifications account for the expression level divergence (Fig. 3) between parental and new genes. While their spatiotemporal expression differences (Fig. 2B) are more likely to be caused by a different composition of CREs. To test this, we compared the enhancer repertoire between new and parental genes, which is annotated by STARR-seq (self-transcribing active regulatory region sequencing) in five Drosophila species(Arnold et al., 2014) or by CAGE (cap analysis of gene expression) in human(Andersson et al., 2014). These enhancers show an expected enrichment (H3K4me1, H3K27ac) or depletion (H3K27me3) of specific histone modifications (**Supplementary Fig. 20**). We found that enhancers of Drosophila new genes show a significant (*P*<0.01, Wilcoxon test) lower level of H3K27ac modifications compared to those of parental genes (**Supplementary Fig. 21**), which probably results in a lower expression level of some new genes. New genes exhibit gains, losses, or sequence mutations of their enhancers compared to those of their parental genes. And a higher percentage of new genes have undergone such turnovers, but have shown rare retentions of parental genes’ enhancers (‘enhancer duplication’, Fig. 5A-B) in older age groups in both Drosophila and human. This indicates a much more diverged cis-regulatory circuit between parental and new genes over evolution. In particular, when searching for the orthologous sequences of specific enhancers (‘enhancer gain’) that were gained by Drosophila new genes in their outgroup species, we found that they are also predominantly enhancers, suggesting new genes have frequently recruited pre-existing enhancers as their new CREs (Fig. 5C). In fact, 37 out of 173 analysed Drosophila new gene specific enhancers that do not have an ortholog in outgroup species all belong to *de novo* genes. This raises the interesting question about the role of *de novo* enhancers during the emergence of *de novo* genes.

Besides the numbers of enhancers, new genes and parental genes have also diverged for their types of enhancers. It has been recently shown that *D. melanogaster* enhancers can be divided into two classes, according to their specificity to core promoters of either housekeeping genes or developmentally regulated genes, with each enriched for separate classes of sequence motifs(Zabidi et al., 2015). In parallel, human enhancers have also been annotated to have ubiquitous or cell-type/tissue specific expression(Andersson et al., 2014). We compared these two types of enhancers’ distributions between new and parental genes in both human and Drosophila. Indeed, for new genes produced by DNA-based gene duplication, there are significantly (*P*<0.05, t-test) fewer housekeeping/ubiquitous enhancers in new genes than parental genes. While retrogenes possess more (*P*>0.05, t-test) developmental/tissue-specific enhancers than their parental genes (Fig. 5D). Correspondingly, there are also fewer housekeeping gene related sequence motifs (e.g., Ohler motif 7) in new DNA-based gene duplicates, while there are more tissue-specific gene related sequence elements (e.g., TATA box and Initiator element) among new retrogenes, comparing to their parental genes (**Supplementary Fig. 22**). These results together demonstrated that the CREs have become diverged for their numbers and types between new and parental genes, which underlies their observed different expression patterns.

## Discussion

Genome-wide and experimental case studies have demonstrated that functional new genes have frequently emerged during evolution and constitute a main driving force underlying the evolution of organismal complexity(S. Chen et al., 2013; Kaessmann, 2010). Similar to any other types of mutations, a nascent gene does not usually confer an immediate selective advantage that will drive its rapid fixation throughout the population. Many of them seem to be initially segregating within the population, and to start out as testis-specific genes. This has been observed for new gene cases of Drosophila, human(Kaessmann, 2010) and recently those that are preferentially expressed in pollen of rice and *Arabidopsis thaliana*(Cui et al., 2015), suggesting male reproductive tissues are a universal cradle for the birth of new genes. Importantly, we further uncovered here an age-dependent transition of new genes from testis-specific genes to become broadly expressed housekeeping genes, supporting the ‘out of testis’ hypothesis that some new genes will evolve much more important functions beyond testis during later evolution. Several factors probably account for such a preference of male reproductive tissues: first, testes of Drosophila and mammals have a distinct epigenomic regulation program from other somatic tissues which may license more promiscuous transcription. This affords the prerequisite of exposing the nascent gene to natural selection. Particularly in mammals, RNA polymerase II is enriched in testis(Schmidt & Schibler, 1995). And recently a testis-specific histone H3 variant H3t that is essential for spermatogenesis has been identified to form a flexible open chromatin structure for allowing more transcription(Ueda et al., 2017). While the Drosophila testis does not show a canonical bivalent chromatin domain with both H3K27me3 and H3K4me3 on differentiation genes as regularly observed in somatic tissues(Gan et al., 2010), many testis-specific genes instead reside in ‘BLACK’ chromatin(Filion et al., 2010) associated with lamin(Shevelyov et al., 2009), which suppresses their somatic expression. These regulation programs ensure a more robust expression of nascent genes in testis, and also restrict their potentially harmful expression in other tissues. Indeed, the second factor is probably due to the selection against the redundant gene dosage in somatic tissues posed by the new genes. As a response to such selection, it has been shown in yeast and mammals, that the expression level of duplication gene is reduced to maintain the gene dosage(Qian, Liao, Chang, & Zhang, 2010). And the selection against a new pleiotropic or broadly expressed gene is expected to be much stronger than that of a tissue-specific gene. This probably accounts for the pattern that many new genes, particularly retrogenes tend to emerge from a pre-existing silencing/regulatory H3K27me3 domain (Fig. 3). It is noteworthy that out-of-testis pattern of retrogenes have been recently reported in mammals(Carelli et al., 2016). However, the pattern observed in this work is mainly attributed to the DNA-level gene duplications, because they outnumber retrogenes and *de novo* genes in every age group. Unlike retrogenes, young gene duplicates are very likely to inherit their parental genes’ regulatory elements with few changes (Fig. 5), and thus also the expression pattern. Such a mechanistic factor is reflected by a similar out-of-testis pattern observed for the parental genes of human and Drosophila (Fig. 1).

A following fundamental question is how did new genes acquire novel and important functions beyond the testis? We addressed the underlying regulatory mechanisms by uncovering an age-dependent acquisition of active histone marks and more turnovers of CREs among both Drosophila and human new genes, consistently across somatic tissues and developmental stages (Fig. 4-5). This suggested that the general evolution trajectory of genes involves becoming more active in chromatin configuration, and more complex in cis-regulatory circuits. The change of repressive histone marks, however show variations between species and between somatic and germline tissues. The interspecific difference may be attributed to the presence and absence of DNA methylation in human and Drosophila respectively. It has been reported that the level of promoter DNA methylation, which is negatively associated with gene expression level, also becomes lower in older human gene duplicate pairs(Keller & Soojin, 2014). This indicated that in human, DNA methylation synergistically acts with an age-dependent loss of repressive histone marks and results in more active genes in older age groups observed in this study. However, Drosophila lack DNA methylation except for a very low level of methylation at early embryonic stages(Takayama et al., 2014). Another study recently showed that Drosophila and mouse employ different histone modifications for balancing the gene dosage after gene duplication(Chang & Liao, 2017). These factors, as well as a mixed cell types (late embryos and larvae) used for Drosophila ChIP-seq data probably together account for the different trajectories of repressive histone marks along the age groups between Drosophila and human, for both new and parental genes. It is important to note that the major differences between previous studies(Arthur et al., 2014; Chang & Liao, 2017; Keller & Soojin, 2014) of epigenetic modifications on gene duplications and this work are that the former focused on the comparison between duplicated genes vs. single-copy genes, and there was no distinction between parental and new gene copies. However, as shown here, because parental genes are by definition older than new genes, they can have very different trajectories of epigenetic changes (Fig. 4).

Finally, we uncovered that parental and new genes have clearly diverged for their CRE repertoire and become enriched for different types of enhancers or sequence motifs. Despite the much progress that has been made in identifying the enhancers in a high-throughput manner(Andersson et al., 2014; Arnold et al., 2014), assigning them to their downstream genes still remains a great challenge. We conservatively restricted our analyses to enhancers and their nearby genes in this study, which is an underestimate of the CREs. Comparing to promoters, enhancers seem to have a faster evolution rate(Villar et al., 2015). And a pre-existing enhancer might switch its downstream target to the new gene upon its birth, and facilitate its functional innovation (Fig. 5). It is therefore of great interest in the future to investigate how the numbers and combination of enhancers evolved across different ages of new genes, when more data (e.g., Hi-C) becomes available. As studying new genes’ evolution throughout their life history provides an entry point into understanding the evolution trajectory of genes in general.

## Methods

### Inferring age and origination mechanisms of new genes

We adopted a whole genome alignment based pipeline to identify the new genes and infer their origination time and mechanisms, as described in(Y. E. Zhang et al., 2010). For *Drosophila* (Ensembl metazoa release 26) and human (Ensembl release v73), we took advantage of UCSC whole genome syntenic alignment (https://genome.ucsc.edu/cgi-bin/hgGateway) and inferred the phylogenetic distribution of orthologs of *D. melanogaster* or human genes among the other Drosophila or vertebrate genomes. Species or lineage-specific genes were identified as new genes and then assigned into respective age groups, based on their presence/absence in outgroup species and parsimony. We classified new genes’ origination mechanisms as DNA-based duplication (gene duplication), RNA-based duplication (retroposition) and *de novo* origination. We characterized retrogenes as those intronless genes whose parental genes have at least one intron. Otherwise, it will be classified as gene duplication. A gene will be defined as *de novo* gene if no alignment hit can be found in the outgroup protein repertories with a BLAST(Camacho et al., 2009) e-value cut-off as 10^−6^, an alignment length cut-off as 70%, and a sequence identity cut-off as 50%, also without any annotated paralogs by Ensembl.

### Transcriptomic and epigenomic analyses

To differentiate between the parental and new gene sequences, we used MUSCLE (v.3.8.31)(Y. E. Zhang et al., 2010) and produced pairwise alignments for 452 Drosophila and 1351 human parental-new gene pairs. Using parental genes as a reference, we recorded the nucleotide and genomic position information of all diagnostic SNPs between new and parental genes, with customized perl codes. Transcriptomic and epigenomic data of *D. melanogaster* and human were retrieved from databases of NCBI (https://www.ncbi.nlm.nih.gov/sra), ENCODE (https://www.encodeproject.org/), and Roadmap Epigenomics project (http://www.roadmapepigenomics.org/), and published single-cell sequencing data(Xue et al., 2013; Yan et al., 2013) (for all data resources: **Supplementary Table 2**). We mapped the RNA-seq reads with HISAT2(v2.0.5)(Kim, Langmead, & Salzberg, 2015), and ChIP-seq reads with Bowtie2(v2.2.9)(Langmead & Salzberg, 2012) to the reference genomes of *D. melanogaster* (r6.02) and human (hg19), using a mapping quality cut-off of 20 and taking paired-end relationship into account. We counted the number of RNA-seq or ChIP-seq reads that span the diagnostic SNPs after assigning them to either parental or new genes based on their matched nucleotides. For *de novo* genes without any parental genes, we used BEDTools (v2.25.0)(Quinlan & Hall, 2010) to count the total read number within the gene regions. After calibrating the difference of total sequenced reads between different samples, the RNA-seq read number of each gene was then normalized against the corresponding genomic DNA-seq read number to correct for the mapping bias, and also allow for comparison between genes. Similarly, we calculated the log2 ratio of ChIP (IP) vs. input (IN) reads that span the diagnostic SNPs, for the entire gene region for the markers H3K36me1, H3K36me3, H3K9me2, H3K9me3, H3K27me3, H3K79me1, H3K79me2, H4K16ac; or specifically for the putative promoter region (+/− 2kb around the transcriptional start sites) for the markers H3K4me1, H3K4me2, H3K4me3, H3K18ac, H3K27ac, H3K9ac and RNA polymerase II. To test for the validity of normalization, we performed correlation analyses between the gene expression level vs. the histone modification level with R, which showed consistent results with those derived from ENCODE or modENCODE project. We defined a gene as being transcriptionally active or bound by certain histone modifications, based on the distribution of expression level of normalized histone modification level of all genes or promoters in the respective tissue or stage. Bivalent genes were defined as those bound by both H3K27me3 and H3K4me3.

We used two Drosophila RNA-FISH databases, Fly-FISH(Lécuyer et al., 2007) and Berkeley Drosophila Genome Project (BDGP)(Tomancak et al., 2002) for comparing the localization patterns between new and parental genes during embryogenesis. We combined the two databases and when there were overlapping genes between the two, we selected genes with their parental or new gene’s data available in the same database. Then we compared the annotated anatomical terms and embryonic stages with detected expression between new and parental genes.

### CRE data analysis

We used a non-redundant enhancer dataset annotated for *D. melanogaster, D. yakuba, D. ananassae, D. pseudoobscura* and *D. willistoni* by STARR-seq(Arnold et al., 2014), and a human enhancer dataset annotated by FANTOM5 consortium by CAGE(Andersson et al., 2014). For the Drosophila housekeeping or developmental gene enhancers, we used the data from(Zabidi et al., 2015), and for human ubiquitous or tissue/cell-specific enhancer, we used the data from FANTOM5 consortium(Andersson et al., 2014). We assigned the enhancer-gene relationship following the same rules as the published work: for STARR-seq enhancers, they are assigned to either parental or new genes when they fall within 2kb up- or down-stream of the TSS; for the FANTOM5 enhancers, they are assigned to either parental or new genes when they fall within 5kb up- or down-stream of the TSS. For the Drosophila housekeeping/developmental gene enhancers, we additionally include those that located within the 5 kb upstream from the TSS, within the gene body itself, 2 kb downstream of the gene, as well as the ‘closest enhancer’ which is assigned to the closest TSS of an annotated gene. To verify the enhancer activities, we calculated the log2 transformed IP/IN ratios at the enhancer regions, after aligning the ChIP-seq reads of H3K27me3, H3K4me1 and H3K27ac derived from Drosophila S2 cells and human K562 cells to the respective reference genomes using bowtie2. We defined a ‘enhancer gain’ event when the new gene has a specific enhancer that is absent in the parental gene and also outgroup species (see below), and vice versa for ‘enhancer loss’; while ‘enhancer duplication’ is defined as the case that new and parental genes share the identical enhancer sequence; and ‘enhancer mutation’ refers to the case that new and parental genes have sequence divergences between a pair of homologous enhancers. We examined the candidate cases of enhancer gain/loss using genome alignments between the focal and outgroup species. Once the coordinates of the focal enhancer were translated into those in the outgroup by the UCSC liftOver tool, we further used BEDTools to check the presence/absence of orthologous sequence. For Drosophila, we investigated branches E to B, where STARR-seq annotated enhancers are available for the included species. When examining the source of enhancer gain, if the orthologous sequence of the focal enhancer in the outgroup has also been annotated as an enhancer, we defined the gained enhancer as a pre-existing enhancer. Otherwise, it is defined as a *de novo* enhancer. We used MEME suite (Bailey et al., 2009) to search for the motif occurrences in the new and parental genes, with the published motif matrixes(Zabidi et al., 2015) as queries.

## Acknowledgement

We are very grateful to Yong Zhang, Da-qi Yu, Chun-yan Chen for providing the new gene datasets, and James Howie for his helpful comments. This project is supported by National Natural Science Foundation of China (31722050, 31671319), the Thousand Talents Plan, European Research Council (grant agreement 677696), the Fundamental Research Funds for the Central Universities, and start-up funds from Zhejiang University to Z.Q.

## Author Contributions

Z.Q. conceived the project, Z.J. and Z.Q. performed the analyses and wrote the manuscript.

## Supplementary Material Legend

**Supplementary Table 1 Percentage of putative pseudogenes across different age groups**

We defined “putative pseudogenes” as gene which show no expression across all investigated developmental stages or tissues. denovo_pse: putative pseudogenes among *de novo*; denovo_all: all the annotated *de novo* genes; dup_new_pse: putative pseudogenes of new DNA-based duplicated genes; dup_new_all: all the DNA-based duplicated new genes.

**Supplementary Table 2 Type and accession numbers of analyzed datasets** Transcriptomic and epigenomic data of *Drosophila melanogaster* and human were retrieved from databases of NCBI (https://www.ncbi.nlm.nih.gov/sra), ENCODE (https://www.encodeproject.org/), and Roadmap Epigenomics project (http://www.roadmapepigenomics.org/). Additional single-cell RNA-seq data of human early embryo and PGC/somatic cells were retrieved from published paper.

**Supplementary Figure 1 Transcriptomic and epigenomic data used in this work** Available and analyzed data are marked with grey color. Drosophila sample abbreviations: E0-4h: embryos within 0-4 hours after egg lay; mc8-14: mitotic cycle 8-14; L1, L2, L3: the first/second/third instar larvae; AF: adult female, AM: adult male; S2: Schneider 2 cells. Human sample abbreviations: EF: embryo female, EM: embryo male; FeF: fetal female, FeM: fetal male; CF: child female, CM: child male; AF: adult female, AM: adult male.

**Supplementary Figure 2 Densities of informative sites between parental and new genes**

We showed SNP densities (SNPs per bp length) for each age group of parental-new gene pairs, for (A) *Drosophila* promoter region;(B) *Drosophila* gene body region; (C) human promoter region;(D) human gene body region

**Supplementary Figure 3 Correlations between ChIP-seq and RNA-seq data**

We performed Pearson’s correlation test between the normalized gene expression level and the histone modification level across all studied Drosophila and human tissues/stages. And we showed the −log10 based *P*-values of the correlation tests, with blank cells for those do not have a significant correlation (*P*>0.05), and crossed cells for those without available data.

Drosophila sample abbreviations: E0-4h: embryos within 0-4 hours after egg lay; mc8-14: mitotic cycle 8-14; L1, L2, L3: the first/second/third instar larvae; AF: adult female, AM: adult male; S2: Schneider 2 cells. Human sample abbreviations: EF: embryo female, EM: embryo male; FeF: fetal female, FeM: fetal male; CF: child female, CM: child male; AF: adult female, AM: adult male.

**Supplementary Figure 4 Percentage of genes bound by histone modifications** We defined bound genes of as those with log2 based ChIP vs. input ratio, or RNA-seq vs. DNA-seq ratio higher than 0, and then we calculated their percentage for each tissue or stage. Blank cells are those without data available.

**Supplementary Figure 5 Distribution of coefficients of expression variation**

We show the density plot with the coefficients of variation of expression levels for *Drosophila* and human genes. The *Drosophila* gene expression levels are calculated based on modENCODE data, and human gene expression levels are calculated based on GTEx data. Based on the entire distribution, we artificially defined genes as stable genes, middle genes and regulated genes, separated by the dotted lines as cutoffs.

**Supplementary Figure 6 Comparison of new and parental genes’ expression levels across tissues and stages**

We show here the percentage of active genes (left columns) and media values of expression levels (right columns) for *Drosophila* and human across different tissues and stages. We compared the expression levels between new and parental genes with t-test, and mark those with significant differences with asterisks (*: *P* < 0.05; **: *P* < 0.01; ***: *P* <0.001). If the entire column is significant, we show the asterisks above the column.

**Supplementary Figure 7 Gene Ontology (GO) enrichment analysis for human zygotically activated retrogenes**

We defined the zygotically activated genes as those which showed a 2-fold increase of expression level at embryonic 8-cell stage, comparing to the 4-cell stage. Then we showed the *P*-values of enriched GO terms based on analyses with DAVID v6.8.

**Supplementary Figure 8 RNA-FISH data with parental genes ubiquitously expressed, and new genes specifically expressed**

Gene expression patterns during *Drosophila* embryogenesis were extracted from BDGP & Fly-FISH databases, with each two columns showing the pattern of new gene (left) and parental gene (right), and each row for the investigated stage.

**Supplementary Figure 9 RNA-FISH data with parental genes specifically expressed, and new genes ubiquitously expressed**

**Supplementary Figure 10 RNA-FISH data with parental genes and new genes specifically expressed at different positions**

**Supplementary Figure 11 Percentage of active genes across different tissues and age groups**

We show the percentage of active genes across each age group (each row) and tissue/stage sample (each column), with *de novo* genes, new genes (gene duplicates or retrogenes) and their parental genes separated for Drosophila (A), human embryonic stages (B), and human tissues (C).

**Supplementary Figure 12 Expression level across different tissues and age groups**

We show here the median expression levels across each age group (each row) and tissue/stage sample (each column), with *de novo* genes, new genes (gene duplicates or retrogenes) and their parental genes separated for Drosophila (A), human embryonic stages (B), and human tissues (C).

**Supplementary Figure 13 New and parental genes show divergent histone modifications**

We show here the distributions of normalised histone modifications (log2 based IP/IN ratio) across different tissues or stages of Drosophila and human. New genes were shown in different colours of solid lines, while parental genes and genome-wide average distribution were shown in coloured areas. We also showed the *P-values* of significance tests comparing the new vs. parental genes (t-test), or the de novo genes vs. genome-wide average (Wilcoxon test).

**Supplementary Figure 14 New genes and parental genes show different levels of histone modifications**

We compared the normalised histone modification levels between parental and new genes across age groups, using *Drosophila* mix-adult sample and human adult male heart left ventricle sample for illustration.

**Supplementary Figure 15 Comparing surrounding genes of new and parental genes for their H3K27me3 histone modification**

We plotted the distributions of H3K27me3 modification levels of the surrounding genes of new genes (in coloured lines) vs. those of parental genes (in blue area), and also those of *de novo* genes vs. all the genes in the genome (in green area).

**Supplementary Figure 16 Bivalent genes of *Drosophila***

We defined the bivalent gene as those bound by both H3K4me3 and H3K27me3, and then we showed the percentage of bivalent genes across *Drosophila* developmental stages.

**Supplementary Figure 17 Bivalent genes of human**

**Supplementary Figure 18 Percentage of bound genes by histone modifications across different tissues and age groups**

We show the percentage of bound genes by respective histone modification marks across different Drosophila (A) developmental stages or human (B) tissues, and age groups (each row).

**Supplementary Figure 19 Levels of histone modifications across different tissues and age groups**

We show the median values of histone modification levels across different Drosophila (A) developmental stages or human (B) tissues, and age groups (each row).

**Supplementary Figure 20 Histone modifications at human and Drosophila enhancer regions**

*Drosophila* enhancers were derived from, human enhancers were from FANTOM5 project. We used ChIP-seq data of Drosophila S2 cells and human k562 cells and produced the metagene plots surrounding the enhancer regions.

**Supplementary Figure. 21 Comparing histone modifications of enhancers between new and parental genes**

Y-axis shows the normalised histone modification levels of new (red) and parental (blue) enhancers.

**Supplementary Figure. 22 Enrichment of sequence motifs in new and parental genes**

*Drosophila* core-promoter motifs are divided as housekeeping gene motifs or developmental gene motifs based on annotations of, and similarly for human sequence motifs based on annotation. Then the average densities of these different kinds of motifs were compared between new and parental genes, with t-test P-values shown (*: *P* < 0.05; **: *P* < 0.01; ***: *P* < 0.001).

## References

Andersson, R., Gebhard, C., Miguel-Escalada, I., Hoof, I., Bornholdt, J., Boyd, M., … Sandelin, A. (2014). An atlas of active enhancers across human cell types and tissues. Nature, 507(7493), 455–461. doi:10.1038/nature1278.

Arnold, C. D., Gerlach, D., Spies, D., Matts, J. A., Sytnikova, Y. A., Pagani, M., … Stark, A. (2014). Quantitative genome-wide enhancer activity maps for five Drosophila species show functional enhancer conservation and turnover during cis-regulatory evolution. Nat Genet, 46(7), 685–692. doi:10.1038/ng.300.

Arthur, R. K., Ma, L., Slattery, M., Spokony, R. F., Ostapenko, A., Negre, N., & White, K. P. (2014). Evolution of H3K27me3-marked chromatin is linked to gene expression evolution and to patterns of gene duplication and diversification. Genome Res, 24(7), 1115–1124. doi:10.1101/gr.162008.11.

Assis, R., & Bachtrog, D. (2013). Neofunctionalization of young duplicate genes in Drosophila. Proc Natl Acad Sci U S A, 110(43), 17409–17414. doi:10.1073/pnas.131375911.

Bailey, T. L., Boden, M., Buske, F. A., Frith, M., Grant, C. E., Clementi, L., … Noble, W. S. (2009). MEME SUITE: tools for motif discovery and searching. Nucleic Acids Res, 37(Web Server issue), W202–208. doi:10.1093/nar/gkp33.

Betran, E. (2015). The “life histories” of genes. J Mol Evol, 80(3-4), 186–188. doi:10.1007/s00239-015-9668-.

Braude, P., Bolton, V., & Moore, S. (1988). Human gene expression first occurs between the four-and eight-cell stages of preimplantation development. Nature, 332(6163), 459.

Camacho, C., Coulouris, G., Avagyan, V., Ma, N., Papadopoulos, J., Bealer, K., & Madden, T. L. (2009). BLAST+: architecture and applications. BMC Bioinformatics, 10, 421. doi:10.1186/1471-2105-10-42.

Carelli, F. N., Hayakawa, T., Go, Y., Imai, H., Warnefors, M., & Kaessmann, H. (2016). The life history of retrocopies illuminates the evolution of new mammalian genes. Genome Res, 26(3), 301–314. doi:10.1101/gr.198473.11.

Carvunis, A. R., Rolland, T., Wapinski, I., Calderwood, M. A., Yildirim, M. A., Simonis, N., … Vidal, M. (2012). Proto-genes and de novo gene birth. Nature, 487(7407), 370–374. doi:10.1038/nature1118.

Chang, A. Y., & Liao, B. Y. (2017). Recruitment of histone modifications to assist mRNA dosage maintenance after degeneration of cytosine DNA methylation during animal evolution. Genome Res, 27(9), 1513–1524. doi:10.1101/gr.221739.11.

Chen, S., Krinsky, B. H., & Long, M. (2013). New genes as drivers of phenotypic evolution. Nat Rev Genet, 14(9), 645–660. doi:10.1038/nrg352.

Chen, S., Zhang, Y. E., & Long, M. (2010). New genes in Drosophila quickly become essential. Science.

Consortium, E. P. (2012). An integrated encyclopedia of DNA elements in the human genome. Nature, 489(7414), 57–74. doi:10.1038/nature1124.

Consortium, G., Laboratory, D. A., Coordinating Center-Analysis Working, G., Statistical Methods groups-Analysis Working, G., Enhancing, G. g., Fund, N. I. H. C., … Montgomery, S. B. (2017). Genetic effects on gene expression across human tissues. Nature, 550(7675), 204–213. doi:10.1038/nature2427.

Consortium, m., Roy, S., Ernst, J., Kharchenko, P. V., Kheradpour, P., Negre, N., … Kellis, M. (2010). Identification of functional elements and regulatory circuits by Drosophila modENCODE. Science, 330(6012), 1787–1797. doi:10.1126/science.119837.

Cui, X., Lv, Y., Chen, M., Nikoloski, Z., Twell, D., & Zhang, D. (2015). Young genes out of the male: an insight from evolutionary age analysis of the pollen transcriptome. Mol Plant, 8(6), 935–945. doi:10.1016/j.molp.2014.12.00.

Dai, H., Chen, Y., Chen, S., Mao, Q., Kennedy, D., Landback, P., … Long, M. (2008). The evolution of courtship behaviors through the origination of a new gene in Drosophila. Proc Natl Acad Sci U S A, 105(21), 7478–7483. doi:10.1073/pnas.080069310.

Ding, Y., Zhao, L., Yang, S., Jiang, Y., Chen, Y., Zhao, R., … Wang, W. (2010). A young Drosophila duplicate gene plays essential roles in spermatogenesis by regulating several Y-linked male fertility genes. PLoS Genet, 6(12), e1001255. doi:10.1371/journal.pgen.100125.

Domazet-Loso, T., & Tautz, D. (2010). A phylogenetically based transcriptome age index mirrors ontogenetic divergence patterns. Nature, 468(7325), 815–818. doi:10.1038/nature0963.

Filion, G. J., van Bemmel, J. G., Braunschweig, U., Talhout, W., Kind, J., Ward, L. D., … van Steensel, B. (2010). Systematic protein location mapping reveals five principal chromatin types in Drosophila cells. Cell, 143(2), 212–224. doi:10.1016/j.cell.2010.09.00.

Gan, Q., Schones, D. E., Eun, S. H., Wei, G., Cui, K., Zhao, K., & Chen, X. (2010). Monovalent and unpoised status of most genes in undifferentiated cell-enriched Drosophila testis. Genome Biol, 11(4), R42.

Guschanski, K., Warnefors, M., & Kaessmann, H. (2017). The evolution of duplicate gene expression in mammalian organs. Genome Res, 27(9), 1461–1474. doi:10.1101/gr.215566.11.

Ho, J. W., Jung, Y. L., Liu, T., Alver, B. H., Lee, S., Ikegami, K., … Park, P. J. (2014). Comparative analysis of metazoan chromatin organization. Nature, 512(7515), 449–452. doi:10.1038/nature1341.

Kaessmann, H. (2010). Origins, evolution, and phenotypic impact of new genes. Genome Res, 20(10), 1313–1326. doi:10.1101/gr.101386.10.

Kalinka, A. T., Varga, K. M., Gerrard, D. T., Preibisch, S., Corcoran, D. L., Jarrells, J., … Tomancak, P. (2010). Gene expression divergence recapitulates the developmental hourglass model. Nature, 468(7325), 811–814. doi:10.1038/nature0963.

Katju, V., & Lynch, M. (2006). On the formation of novel genes by duplication in the Caenorhabditis elegans genome. Mol Biol Evol, 23(5), 1056–1067. doi:10.1093/molbev/msj11.

Keller, T. E., & Soojin, V. Y. (2014). DNA methylation and evolution of duplicate genes. Proc Natl Acad Sci U S A, 111(16), 5932–5937.

Kharchenko, P. V., Alekseyenko, A. A., Schwartz, Y. B., Minoda, A., Riddle, N. C., Ernst, J., … Park, P. J. (2011). Comprehensive analysis of the chromatin landscape in Drosophila melanogaster. Nature, 471(7339), 480–485. doi:10.1038/nature0972.

Kim, D., Langmead, B., & Salzberg, S. L. (2015). HISAT: a fast spliced aligner with low memory requirements. Nat Methods, 12(4), 357–360. doi:10.1038/nmeth.331.

Knowles, D. G., & McLysaght, A. (2009). Recent de novo origin of human protein-coding genes. Genome Res, 19(10), 1752–1759. doi:10.1101/gr.095026.10.

Kondo, S., Vedanayagam, J., Mohammed, J., Eizadshenass, S., Kan, L. J., Pang, N., … Lai, E. C. (2017). New genes often acquire male-specific functions but rarely become essential in Drosophila. Genes Dev, 31(18), 1841–1846. doi:10.1101/gad.303131.11.

Langmead, B., & Salzberg, S. L. (2012). Fast gapped-read alignment with Bowtie 2. Nat Methods, 9(4), 357–359. doi:10.1038/nmeth.192.

Lécuyer, E., Yoshida, H., Parthasarathy, N., Alm, C., Babak, T., Cerovina, T., … Krause, H. M. (2007). Global analysis of mRNA localization reveals a prominent role in organizing cellular architecture and function. Cell, 131(1), 174–187.

Lesch, B. J., Silber, S. J., McCarrey, J. R., & Page, D. C. (2016). Parallel evolution of male germline epigenetic poising and somatic development in animals. Nat Genet, 48(8), 888–894. doi:10.1038/ng.359.

Long, M., & Langley, C. H. (1993). Natural selection and the origin of jingwei, a chimeric processed functional gene in Drosophila. Science.

Luis Villanueva-Canas, J., Ruiz-Orera, J., Agea, M. I., Gallo, M., Andreu, D., & Alba, M. M. (2017). New genes and functional innovation in mammals. Genome Biol Evol, 9(7), 1886–1900. doi:10.1093/gbe/evx13.

Lynch, M., & Force, A. (2000). The probability of duplicate gene preservation by subfunctionalization. Genetics, 154(1), 459–473.

Ohno, S. (1970). Evolution by gene duplication: Springer, Berlin, Heidelberg.

Palmieri, N., Kosiol, C., & Schlotterer, C. (2014). The life cycle of Drosophila orphan genes. Elife, 3, e01311. doi:10.7554/eLife.0131.

Perez-Lluch, S., Blanco, E., Tilgner, H., Curado, J., Ruiz-Romero, M., Corominas, M., & Guigo, R. (2015). Absence of canonical marks of active chromatin in developmentally regulated genes. Nat Genet, 47(10), 1158–1167. doi:10.1038/ng.338.

Qian, W., Liao, B.-Y., Chang, A. Y.-F., & Zhang, J. (2010). Maintenance of duplicate genes and their functional redundancy by reduced expression. Trends Genet, 26(10), 425–430.

Quinlan, A. R., & Hall, I. M. (2010). BEDTools: a flexible suite of utilities for comparing genomic features. Bioinformatics, 26(6), 841–842. doi:10.1093/bioinformatics/btq03.

Ross, B. D., Rosin, L., Thomae, A. W., Hiatt, M. A., Vermaak, D., de la Cruz, A. F., … Malik, H. S. (2013). Stepwise evolution of essential centromere function in a Drosophila neogene. Science, 340(6137), 1211–1214. doi:10.1126/science.123439.

Ruiz-Orera, J., Hernandez-Rodriguez, J., Chiva, C., Sabido, E., Kondova, I., Bontrop, R., … Alba, M. M. (2015). Origins of de novo genes in human and chimpanzee. PLoS Genet, 11(12), e1005721. doi:10.1371/journal.pgen.100572.

Schmidt, E. E., & Schibler, U. (1995). High accumulation of components of the RNA polymerase II transcription machinery in rodent spermatids. Development, 121(8), 2373–2383.

Shevelyov, Y. Y., Lavrov, S. A., Mikhaylova, L. M., Nurminsky, I. D., Kulathinal, R. J., Egorova, K. S., … Nurminsky, D. I. (2009). The B-type lamin is required for somatic repression of testis-specific gene clusters. Proc Natl Acad Sci U S A, 106(9), 3282–3287. doi:10.1073/pnas.081193310.

Sysoev, V. O., Fischer, B., Frese, C. K., Gupta, I., Krijgsveld, J., Hentze, M. W., … Ephrussi, A. (2016). Global changes of the RNA-bound proteome during the maternal-to-zygotic transition in Drosophila. Nat Commun, 7, 12128. doi:10.1038/ncomms1212.

Takayama, S., Dhahbi, J., Roberts, A., Mao, G., Heo, S. J., Pachter, L., … Boffelli, D. (2014). Genome methylation in D. melanogaster is found at specific short motifs and is independent of DNMT2 activity. Genome Res, 24(5), 821–830. doi:10.1101/gr.162412.11.

Tomancak, P., Beaton, A., Weiszmann, R., Kwan, E., Shu, S., Lewis, S. E., … Rubin, G. M. (2002). Systematic determination of patterns of gene expression during Drosophila embryogenesis. Genome Biol, 3(12), RESEARCH0088.

Ueda, J., Harada, A., Urahama, T., Machida, S., Maehara, K., Hada, M., … Osakabe, A. (2017). Testis-specific histone variant H3t gene is essential for entry into spermatogenesis. Cell reports, 18(3), 593–600.

Villar, D., Berthelot, C., Aldridge, S., Rayner, T. F., Lukk, M., Pignatelli, M., … Odom, D. T. (2015). Enhancer evolution across 20 mammalian species. Cell, 160(3), 554–566. doi:10.1016/j.cell.2015.01.00.

Wu, D. D., Irwin, D. M., & Zhang, Y. P. (2011). De novo origin of human protein-coding genes. PLoS Genet, 7(11), e1002379. doi:10.1371/journal.pgen.100237.

Xue, Z., Huang, K., Cai, C., Cai, L., Jiang, C. Y., Feng, Y., … Fan, G. (2013). Genetic programs in human and mouse early embryos revealed by single-cell RNA sequencing. Nature, 500(7464), 593–597. doi:10.1038/nature1236.

Yan, L., Yang, M., Guo, H., Yang, L., Wu, J., Li, R., … Tang, F. (2013). Single-cell RNA-Seq profiling of human preimplantation embryos and embryonic stem cells. Nat Struct Mol Biol, 20(9), 1131–1139. doi:10.1038/nsmb.266.

Zabidi, M. A., Arnold, C. D., Schernhuber, K., Pagani, M., Rath, M., Frank, O., & Stark, A. (2015). Enhancer-core-promoter specificity separates developmental and housekeeping gene regulation. Nature, 518(7540), 556–559. doi:10.1038/nature1399.

Zhang, W., Landback, P., Gschwend, A. R., Shen, B., & Long, M. (2015). New genes drive the evolution of gene interaction networks in the human and mouse genomes. Genome Biol, 16, 202. doi:10.1186/s13059-015-0772-.

Zhang, Y. E., Vibranovski, M. D., Landback, P., Marais, G. A., & Long, M. (2010). Chromosomal redistribution of male-biased genes in mammalian evolution with two bursts of gene gain on the X chromosome. PLoS Biol, 8(10). doi:10.1371/journal.pbio.100049.

Zhao, L., Saelao, P., Jones, C. D., & Begun, D. J. (2014). Origin and spread of de novo genes in Drosophila melanogaster populations. Science.

Zhou, Q., Zhang, G., Zhang, Y., Xu, S., Zhao, R., Zhan, Z., … Wang, W. (2008). On the origin of new genes in Drosophila. Genome Res, 18(9), 1446–1455. doi:10.1101/gr.076588.10.

